# Altered protein quality control contributes to noise-induced hearing loss

**DOI:** 10.1101/452698

**Authors:** Nopporn Jongkamonwiwat, Ann C. Y. Wong, Miguel A Ramirez, Kwang Pak, Yi-Zhi Wang, Allen F. Ryan, Jeffrey N. Savas

## Abstract

Exposure to damaging levels of noise is the most common cause of hearing loss and impairs high frequency hearing in more than 15 % of adult Americans. Using mice exposed to increasing levels of noise in combination with quantitative proteomics, we tested how noise insults remodel the cochlear proteome both acutely and after a two-week recovery period. We used ABR & DPOAE recordings to define the intensity of noise exposure necessary to produce temporary or permanent threshold shifts (TTS, PTS) in young adult mice and found noise at 94 and 105 dB SPL levels for 30 minutes elicits TTS and PTS, respectively. We quantified thousands of proteins and found that noise insults cause a rapid increase rather than a decrease in the levels of many proteins involved with protein homeostasis, myelin, cytoskeletal structures, and cell junctions such as the synapse. The vast majority of proteins with increased levels immediately after noise exposure showed normal levels after two weeks of recovery. However, several proteins involved in oxidative stress and neuroprotection had significantly increased levels only after the recovery period suggesting they play in important role in regeneration. Interestingly, a small panel of mitochondrial proteins were significantly altered only in PTS conditions suggesting potential discrete protein mechanisms. Our discovery-based proteomic analysis extends the recent description of noise-induced cochlear synaptopathy and shows that noise insults drive a robust proteostasis response. These data provide a new understanding of noise sensitive proteins and may inform the development of effective preventiative strategies or therapies for NIHL.

## Introduction

Noise-induced hearing loss (NIHL) is a major health problem affecting hundreds of millions of people. Exposure to damaging levels of noise occurs through occupational, residential, and recreational activities (1, 2). The major cellular substrates of NIHL are the mechanoreceptive cochlear sensory epithelial hair cells (HCs) within the organ of Corti and their associated auditory afferent fibers (ANF). Intense noise at 115-125 dB SPL causes a direct mechanical destruction of HCs in part by damaging their mechano-sensory stereociliary bundles and physically impairing their ability to transduce auditory information (3, 4). Alternatively, lower levels of injurious noise at 85 - 115 dB SPL induces metabolic changes that damage HCs and ANFs. This is because constant stimulation of the auditory system places excess metabolic stress leading to an increase in oxygen consumption, free radical production, and excitotoxicity (4-6).

Noise-induced damage is categorized based on the duration of hearing impairment. Recovery of normal hearing after acoustic trauma depends on the intensity and duration of the exposure (7). Hearing thresholds elevate immediately after noise exposure and may fully or partially recover after days or weeks. Moderate noise exposure results in a transient attenuation of hearing sensitivity, referred to as a TTS, which decreases auditory sensitivity for a period of days to weeks. PTS are caused by more severe insults and result in irreversible sensorineural hearing loss. In laboratory settings TTS conditions recover after two to four weeks, and any residual threshold elevation after this period are considered permanent (7, 8). The molecular mechanisms and protein networks responsible for threshold elevation and the recovery process observed in TTS are poorly understood.

Intense noise exposure alters synaptic connections between HCs, and ANFs or olivocochlear efferent nerve fibers (9, 10). ANF terminals dramatically swell immediately after noxious noise exposures as a result of glutamatergic excitotoxicity, which can result in decoupling of the pre- and postsynaptic membranes (11). Interestingly, swelling subsides within a few days of exposure and ANFs morphologically recover or regenerate on a similar timeline as auditory thresholds (12). The recovery of the terminals with only minimal ANF death suggests that some neural damage is reversible via regeneration mechanisms in purely TTS conditions (10, 13). However, in TTS conditions, up to half of inner HC ribbon synapses are permanently lost (10, 14). In PTS conditions, the pathological consequences of excessive noise are more severe in nature. This ranges from outer and inner HC death with secondary degeneration of ANFs, to mechanical disruption of the HC mechano-transduction machinery (15). Metabolic changes induced by HC overexposure and excitotoxicity can trigger metabolic decompensation resulting in the swelling of nuclei and mitochondria, as well as cytoplasmic vesiculation (16). Activation of cell stress pathways may lead to apoptosis (17). Acute exposures to noise above 130 dB SPL can cause mechanical destruction leading to the disruption of cell junctions and cell rupture, resulting in the mixing of endolymph and perilymph and potassium toxicity to nearby cells (18, 19).

While there is considerable data regarding the morphological effects of noise damage, we have only a limited molecular understanding of TTS and PTS, and know very little about the protein networks involved in mitigating temporary vs permanent damages in the cochlea. A deep biological understanding of the protein alterations responsible for TTS and PTS, and the recovery mechanisms in TTS conditions may provide new insight towards the therapeutic protection and treatment of NIHL. Mass spectrometry (MS)-based proteomics provides an opportunity to investigate complex biological phenomena by identifying and quantitating thousands of proteins. Previous proteomic studies of the auditory system have provided draft HC and organ of Corti proteomes, and identified protein substrates of chemically induced hearing loss (20-24). Transcriptomic analysis has also been informative and revealed a panel of genes acutely regulated in response to very high noise levels (25). To generate a global analysis of noise effects on cochlear proteins, we applied multiple quantitative proteomic strategies to characterize acute proteome remodeling in PTS and TTS conditions. We found direct and compensatory changes in the levels of discrete proteins. The level of hundreds of non-synaptic proteins are acutely affected by noise exposure and, many proteins had increased levels. In particular, noise exposure triggers a robust increase of many proteostasis proteins including nearly the entire proteasome and many heat shock chaperones. We used orthogonal proteomic experiments to validate 2,281 and 1,831 quantified significant altered proteins in PTS and TTS, respectively. Finally, we performed proteomic measurements two weeks after noise exposure. These experiments identified a small panel of proteins exclusively elevated at this post-exposure time point. The hope is that by identifying protein networks with altered levels after noise exposure we can highlight new targets for future prevention or treatment of NIHL.

## Significance

Multiple quantitative proteomic strategies have determined how damaging auditory stimulation alters the cochlear proteome. Our findings show that moderate and high levels of noise causing temporary and permanent hearing loss drive robust and dose-dependent proteome remodeling. We identified cochlear proteins involved with protein degradation and folding with increased levels after noise exposure suggesting that the proteostasis network plays a key role in NIHL. Defining the changes in the cochlear proteome immediately after noise exposure and during the recovery period has provided a new understanding of the protein networks acutely affected by noise and those involved in the recovery process. Altogether, our findings provide many important protein targets for potential future therapeutic targeting, to prevention or treat NIHL.

## Results

### Hearing loss induced by short-term noise exposure

We set out to identify the protein mechanisms associated with NIHL. To minimize the contribution of age-associated factors we perform our analysis using young adult FVB mice (P50-60). We exposed individual mice to 6-18 kHz octave band noise, at 70, 94, 100, and 105 dB SPL intensities for 30 minutes. ABR tone and click, as well as DPOAE hearing measurements were performed before, in addition to 1, 7, and 14 days after noise exposure. We found that 70 dB SPL exposures represented non-traumatic activation (NTA) of the cochlea and resulted in only a slight increase in click and tone ABR thresholds one day post noise exposure **(*SI Appendix*, Fig. S1 *A*-*B* and S2*A*,)**. DPOAE analysis confirmed that OHC function also recovered to normal levels after one week **(*SI Appendix*, Fig. S1 *C*)**. Wave I amplitude, which indicates strength of synaptic transmission primarily between IHCs and ANFs, was unaffected after 70 dB SPL **(*SI Appendix*, Fig. S1*D*)**. Acoustic overstimulation at 94 dB SPL had similar recovery profiles after seven or 14 days and we observed a near-complete recovery of hearing thresholds after two weeks **(*SI Appendix*, Fig. S1 *E-H* and S2*B*)**. Exposure to 100 dB SPL resulted acutely in highly elevated levels of ABR and DPOAE thresholds as compared to 94 dB SPL. However, after 7 and 14 days of recovery, ABR thresholds recovered almost fully to baseline levels **(*SI Appendix*, Fig. S1*I-K* and S2*C*)**. Exposure to 100 dB SPL was the lowest exposure level tested to show permanent reduction in Wave I amplitudes **(*SI Appendix*, Fig. S1*L*)**. Noise exposure to 105 dB SPL caused severe elevations in threshold levels by DPOAE and ABR to click and tone stimuli. Wave I amplitudes were also significantly reduced, and there was minimal recovery of amplitudes and thresholds after two weeks, indicative of permanent damage **(*SI Appendix*, Fig. S1*M-P* and S2*D*)**. In summary, our findings indicate that 30-minute exposures at 70 dB SPL cause minimal hearing impairments, 94 and 100 dB SPL exposures cause predominantly TTS, while exposure to 105 dB SPL results in a predominantly PTS response.

### Noise exposure alters the level of many cochlear proteins

To investigate how excess noise affects the cochlear proteome in TTS and PTS conditions, we developed a quantitative proteomic strategy using ^15^N-metabolically “heavy” labeled mice. The pooled proteins from multiple ^15^N cochleae facilitate accurate quantification of unlabeled proteins from experimental cochleae, by serving as an internal standard for global proteome quantitation (26). In this way, mice exposed to increasing levels of noise causing NTA, TTS, and PTS remain ^14^N and contain “light” proteins, while unexposed cochleae contain “heavy” proteins. We mixed light and heavy cochlea extracts 1:1, digested the proteins to peptides, and performed multi-dimensional chromatograph with tandem mass spectrometry (MS/MS)-based proteomic analysis (27). To control for potential quantification errors, we used a ratio of ratios analysis paradigm (28), and related each noise exposure condition relative to a group of mice placed in a sound chamber without acoustic exposure (i.e. 0 dB SPL) **(Fig. 1*A*)**. We first focused our attention on proteins with Benjamini-Hochberg adjusted *p*-values < 0.05 (B.H. *p*-value). We performed regression analysis and determined threshold log_2_ fold differences (TLFD) from six control mice comparisons that provided the lowest correlation. The threshold levels of log_2_ fold cut offs were determined at 1.24 and -1.35 for up- and down-regulated fold difference, respectively **(Fig. 1*B*)**. Using this strategy, we determined the number of significantly regulated proteins that increased or decreased in a noise level dependent manner. Overall, many more proteins with increased levels rather than decreased across all levels of noise exposure for 30 minutes. Specifically, we found 21, 115, 226 proteins with significantly increased levels in datasets for NTA, TTS, and PTS, respectively **(Fig. 1*C*, *SI Appendix* and Table S1)**.

**Fig. 1.**
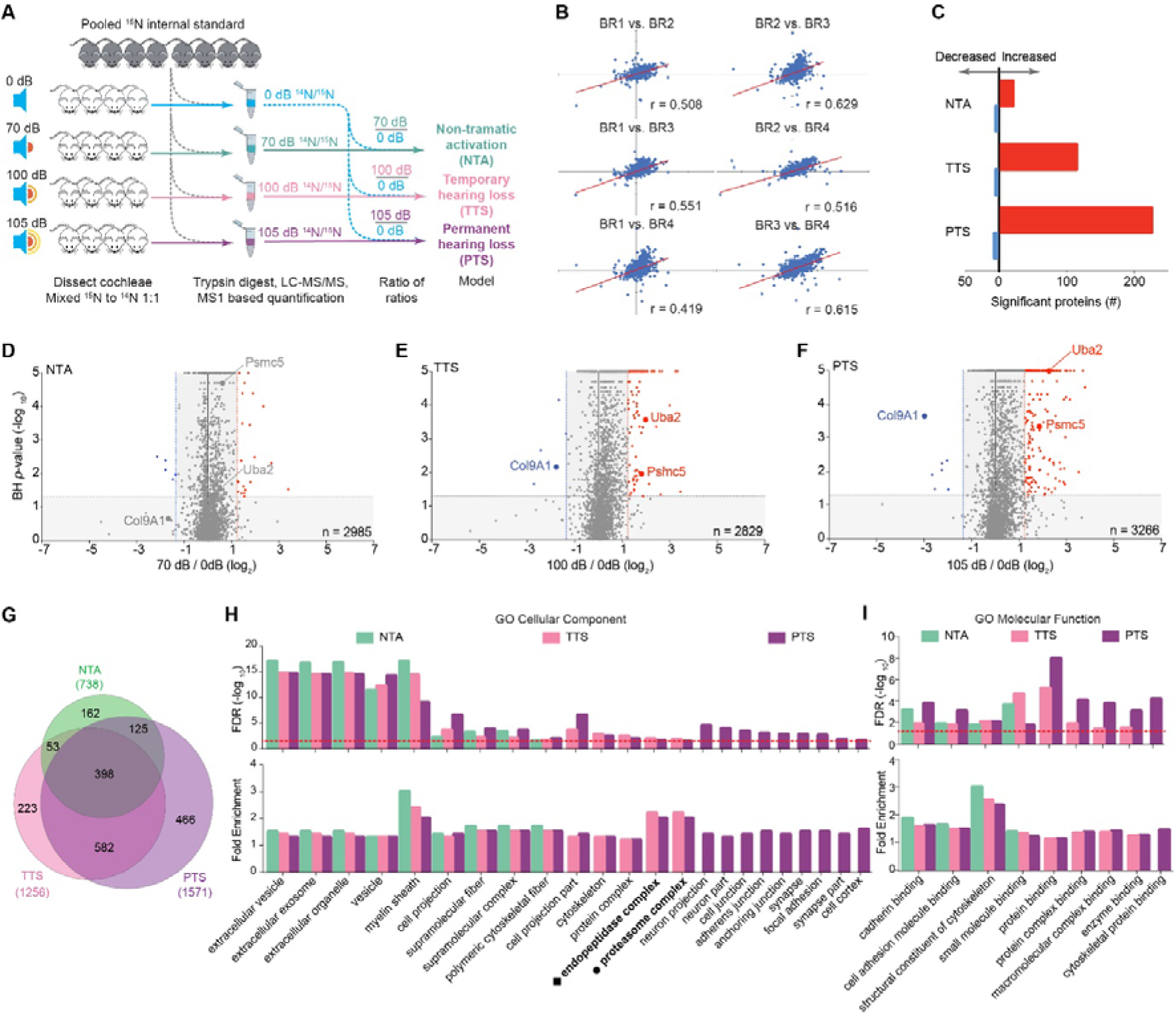
Metabolic stable isotope labeling with MS-based proteomic quantification across noise exposure conditions. (*A*) Experimental scheme for quantifying proteins in different levels of noise exposed cochleae by LC MS/MS using ^15^N labeled cochleae (gray) as an internal standard. (*B*) Correlation plots from pairs of the 4 biological replicates in 0 dB condition were used to verify the level of threshold log fold differences (TLFD) to determine regulated protein expression. (*C*) Summary of total number of significantly quantified proteins which satisfied TLFD criteria. (*D-F*) Volcano plots of the quantified proteins from acoustic overexposed cochleae by LC-MS/MS, graphed as Log_2_ fold change vs. −Log_10_ *P* value. Proteins that satisfied both the statistical cutoff (*P* < 0.05) and TLFD are shown in red or blue points represent up- or down-regulated proteins, respectively. (*G*) Venn diagram of the significantly altered proteins across all three levels of exposure. (H) Enrichment analysis of significant quantified proteins based on GO:Cellular component and, (I) GO: Molecular function terms.

To visualize global trends in cochlear proteome remodeling after increasing levels of noise exposure we graphed our results using volcano plots **(Fig. 1*D-F*)**. In total, our proteomic analysis provided relative quantitation for > 2,800 proteins in each analysis. Interestingly, a panel of proteins (e.g. Psmc5, Uba2) had a noise-dose-dependent increase in their levels after auditory stimulation. A smaller number of proteins (e.g. Col9A1) had noise-dose-dependent decrease in their levels. In total, we identified 668, 1,225 and 1,715 significantly (B.H. *p*-value < 0.05) altered proteins in all three NTA, TTS, and PTS conditions, respectively **(Fig. 1*G*)**. Many proteins were significantly altered in multiple conditions and 356 were significantly altered in all three conditions. To investigate proteome remodeling under less damaging conditions, we performed parallel experiments that limited the duration of noise exposure to 15 min. We again found more proteins with significantly increased levels compared to those with decreased levels **(*SI Appendix*, Fig. S3*A-C* and Table S2)**.

### Noise-increased proteins identify distinct functional processes

To investigate if the significantly altered proteins in the 30 minute datasets (B.H. *p-*value < 0.05) localize to common cellular components or have shared molecular functions we performed ontology cell component (GO: CC) and molecular function (GO:MF) within the PANTHER classification system (29). Proteins altered exclusively in the PTS condition are associated with the GO:CC terms neuronal projections, synapses, cell junctions, among other structures **(Fig. 1*H*)**. Proteins significantly altered in both TTS and PTS conditions, were significantly (Fisher's Exact adjusted FDR < 0.05) enriched for the terms cytoskeleton, cell projection, endopeptidase, and proteasome. These findings support previous evidence that excess noise alters cochlear cell junctions and synapses (10). It is important to note that all GO:CC terms significantly enriched in the NTA dataset (e.g. myelin and cytoskeletal fiber) were also enriched in the TTS and PTS conditions supporting previous findings that TTS and PTS conditions do not simply represent states of enhanced stress but rather distinct biological phenomena **(*SI Appendix*, Table S3)**. Interestingly all of the significantly enriched GO:MF terms are involved with ‘binding’, suggesting that noise exposure even at NTA levels impair many protein-protein interactions **(Fig. 1*H*, *SI Appendix*, Table S3)**.

### Protein alterations across noise exposure intensities

To investigate our datasets further, we homed in on the individual proteins significantly (B.H. *p*-value < 0.05) altered in multiple datasets. Overall, the majority of proteins quantified in more than one condition, had dose dependent increases in abundance with increasing levels of noise **(*SI Appendix*, Table S4)**. Among these noise sensitive proteins, 356 were significantly altered in all three levels of noise exposure, and 603 proteins were significantly altered in both the TTS and PTS datasets. Far fewer proteins were significantly altered in both the NTA and TTS or NTA and PTS conditions **(Fig. 1*G*).** Out of the 398 significant proteins altered in all three conditions, 163 proteins had higher levels in conditions with more intense levels of noise exposure (i.e. PTS > TTS > NTA) **(Fig. 2*A*)**. Similarly, 315 out of the 582 significantly altered proteins quantified in the TTS and PTS conditions had higher levels after exposure to more intense noise (PTS > TTS) **(Fig. 2*B*)**. We also observed a similar pattern of increased protein levels among proteins quantified in the PTS and NTA groups, 79 out of 125 were increased with higher levels of noise (PTS > NTA) **(Fig. 2*C*)**. Finally, we found a similar trend for those proteins significantly altered in the NTA and TTS conditions, 28 out of 53 had higher levels (TTS > NTA). A much smaller panel of proteins had increased reduced levels after intense noise as well **(*SI Appendix*, Fig. S3*A*-*B*)**.

**Fig. 2.**
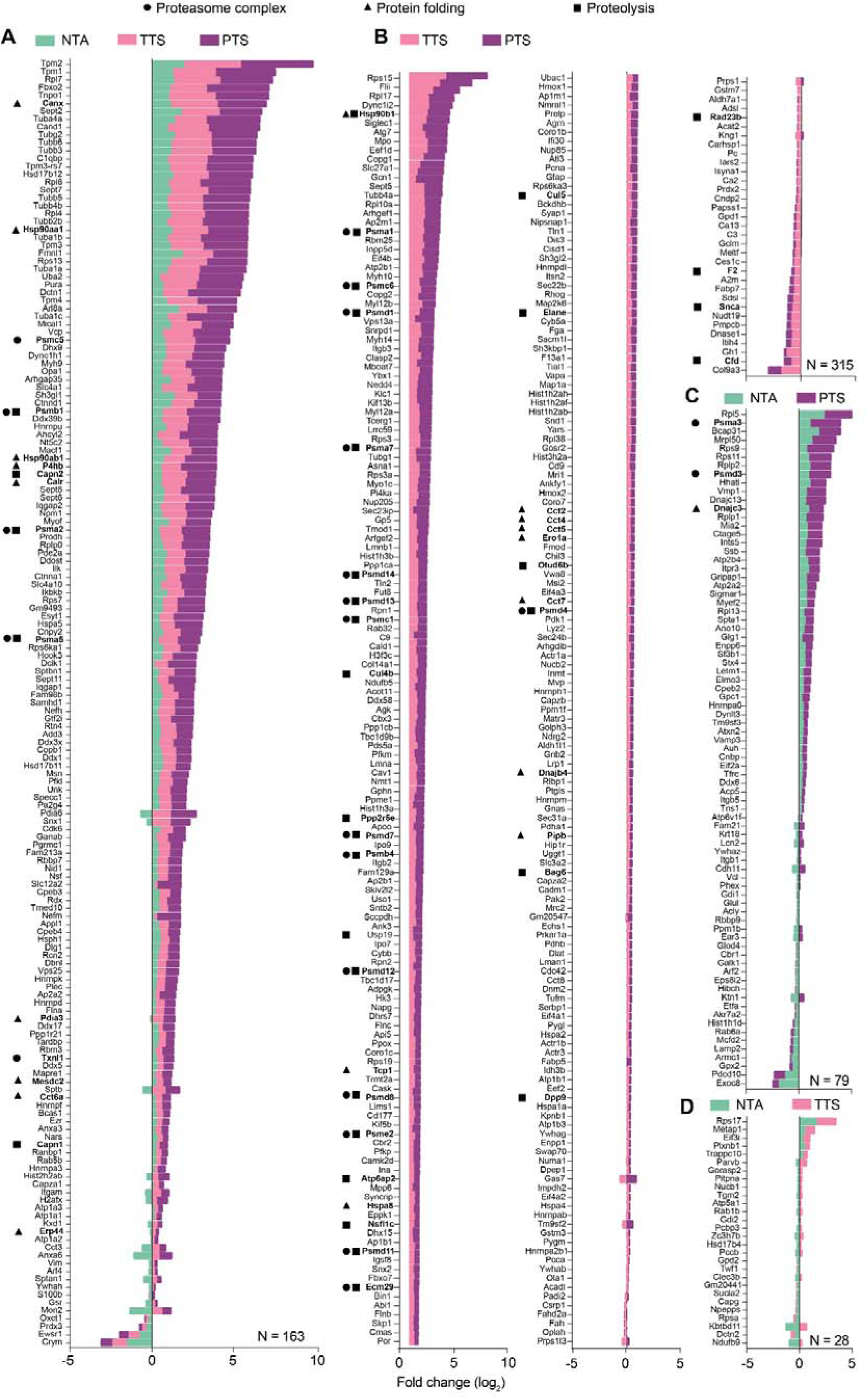
Noise dependent trends of significantly altered proteins across conditions. (*A*) Significant proteins quantified across NTA, TTS and PTS conditions. A majority of these overlapping proteins had increased levels in a noise intensity dependent manner. This phenomena was also observed in groups of proteins measured only in two of the conditions (*B*) PTS:TTS, (*C*) PTS:NTA and (*D*) TTS:NTA. Accumulation of Log_2_ fold change level are presented in the graph together with protein name and the protein function based on GO classification (• = Proteasome complex, ▴ = Protein folding and ▪ = Proteolysis).

Based on our findings that proteasome proteins are significantly enriched in the TTS and PTS but not NTA datasets based on GO:CC analysis **(Fig. 1*H*)**, we searched for proteasome subunits in our datasets of proteins measured in all three levels of noise exposure. Indeed, among those proteins significantly altered in multiple noise exposure conditions, we identified 22 proteasomal proteins, 35 proteins associated with proteolysis and 20 protein-folding factors (based on GO:BP) **(Fig. 2)**. To investigate the possibility that the significantly altered proteins physically interact, we subjected the datasets to STRING analysis. Interestingly, we found additional protein-protein interaction networks and interacting proteins that were significantly increased across noise exposure level (PTS > TTS > NTA). The major protein hubs identified were associated with the proteasome and protein folding (heat shock proteins), suggesting that noise exposure drives a robust proteostasis response **(Fig. 3*A*)**. Noise may drive a protein expression program to delete or refold damaged proteins that could impair cellular functions. Heat shock proteins are involved in various aspects of signal transduction, protein folding, and degradation, apoptosis, and inflammation (30). Two major protein networks were identified in TTS and PTS but not NTA conditions, Arp2/3 complex and the Ubiquinol-Cytochrome C reductase complex **(Fig. 3*B*)**. NADH-Ubiquinone Oxidoreductase (Complex I) was the predominant protein network exclusively altered in the PTS condition which suggests it may contribute specifically to PTS **(Fig. 3*C*)**.

**Fig. 3.**
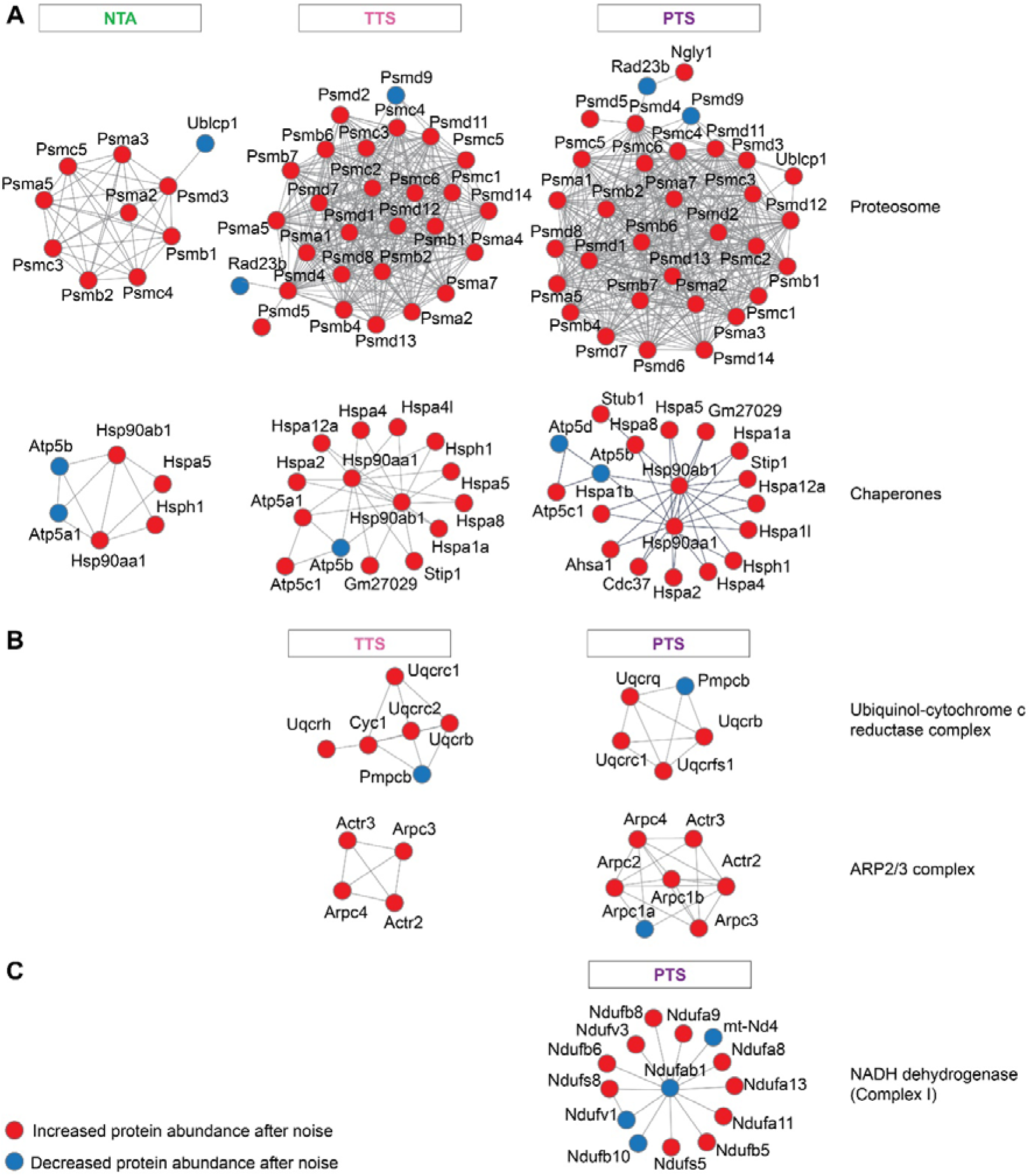
Protein-protein interactions analysis by STRING are represented in five major groups of enzymatic pathways and protein complexes in response to noise stimulation. (*A*) Ubiquitin-proteasome system (UPS) and heat shock protein (HSP) chaperons are the most abundant protein interactions, which also show increased numbers according to the intensity of noise. (*B*) Mitochondrial electron transport complex III enzymes and actin cytoskeletal regulator protein complex ARP2/3 are also shown to increase their ramification in response to higher noise level at TTS and PTS. (*C*) Mitochondrial NADH dehydrogenase (complex I) proteins typically found only in PTS condition which may indicate potentially distinct protein mechanisms for permanent hearing loss.

### Confirmation of ^15^N-based quantitative proteomic results

To confirm our ^15^N-based proteomic measurements we repeated our analysis using 10plex isobaric Tandem mass tags (TMT) which facilitate accurate proteome-wide quantitation (31, 32). Our experimental design consisted of ten animals in three groups: 105 dB SPL (PTS), 94 dB SPL (TTS), and combined 0 and 70 dB SPL (NTA) (***SI Appendix*, Fig. S5*A***). Overall, greater than 70% of proteins quantified with the ^15^N workflow were measured with TMT; 2,179 proteins across both TTS analyses, and 2,401 proteins in both PTS analyses **(*SI Appendix*, Fig. S5*B*-*C*)**. Next, we extracted those proteins in both datasets and compared their levels. In this way, we confirmed protein trends: 757 and 662 up-regulated proteins, and 553 and 556 down-regulated proteins in TTS and PTS conditions, respectively. TMT also confirmed many significant proteins: 90 and 70 proteins with increased levels and 201 and 28 proteins with reduced levels in TTS and PTS conditions **(Fig. S5*D*)**. We also confirmed increased levels of many chaperones in the PTS condition including Bag1, 6, Cct2, 4, 5, 6a, 7 and Clu **(*SI Appendix*, Table S5)**. A representative panel of noise-sensitive proteins show consistent trends of both increased- and decreased-levels across the intensity of noise exposure levels (***SI Appendix,*** Fig. S5*E*, **Table S6)**. The panel of proteins with increased levels included cytoskeletal proteins (Sptan1, Myo6, Tubb4a), a component of the autophagy system (Atg3), nucleopore protein (Nup98), and signaling proteins (Hcls 1). Interestingly, Gephyrin (Gphn) levels were significantly increased in the PTS conditions of both TMT and ^14/15^N datasets. Gphn is a scaffolding molecule that anchors inhibitory neurotransmitter receptors to the postsynaptic cytoskeleton and may reflect an increase in compensatory inhibitory synaptic transmission. Col1a1 had reduced levels presumably due to structural deterioration. Overall, this analysis confirmed the global trend of noise-induced proteome remodeling and many individual protein measurements obtained from the ^15^N quantitative proteomics **(*SI Appendix*, Fig. S5*F*)**.

### Long-term effects of noise exposure

We then explored whether cochlear proteome remodeling was present after a two-week recovery period. Our experimental design consisted of ten animals in three groups 105 (PTS), 94 (TTS), and 70 dB SPL (NTA). Mice were subjected to noise for 30 minutes as previously performed but rather than quantify their proteomes acutely, they were allowed to recover for 14 days, and their cochlear proteome was quantified with TMT-based quantitative proteomics **(Fig. 4*A*)**. More than 76% (n = 1,950) of the proteins measured in the PTS or TTS datasets with ^14^N / ^15^N and over 90% (n = 2,340) from the acute TMT analysis were also measured in the recovery TMT experiment **(Fig. 4*B*)**. We identified a similar number of proteins with altered levels in PTS and TTS conditions in the acute ^14^N / ^15^N or TMT and recovery analyses **(Fig. 4*C*)**. Comparison of the ^14^N / ^15^N acute exposure dataset with the recovery TMT analysis revealed 168 and 109 protein with significantly increased levels,153 and 88 proteins with reduced abundances in PTS and TTS conditions, respectively. Comparisons of the proteins with significantly increased levels between the TMT acute and recovery analyses revealed 17 and 59 proteins in PTS and TTS conditions, respectively. In a similar way, we identified 70 and 250 proteins with significantly reduced levels compared to the PTS and TTS conditions, respectively. Next, we extracted the 114 proteins with significantly altered levels from the acute and recovery TMT analysis. The protein fold change measured in all biological replicates were calculated and represented in heat map **(Fig. 4*D*, *SI Appendix*, Table S7)**. Interestingly, we observed a dramatic change in the protein abundances acutely and after recovery. Hierarchical clustering found 72 and 42 proteins with acutely increased and decreased levels respectively.

**Fig. 4.**
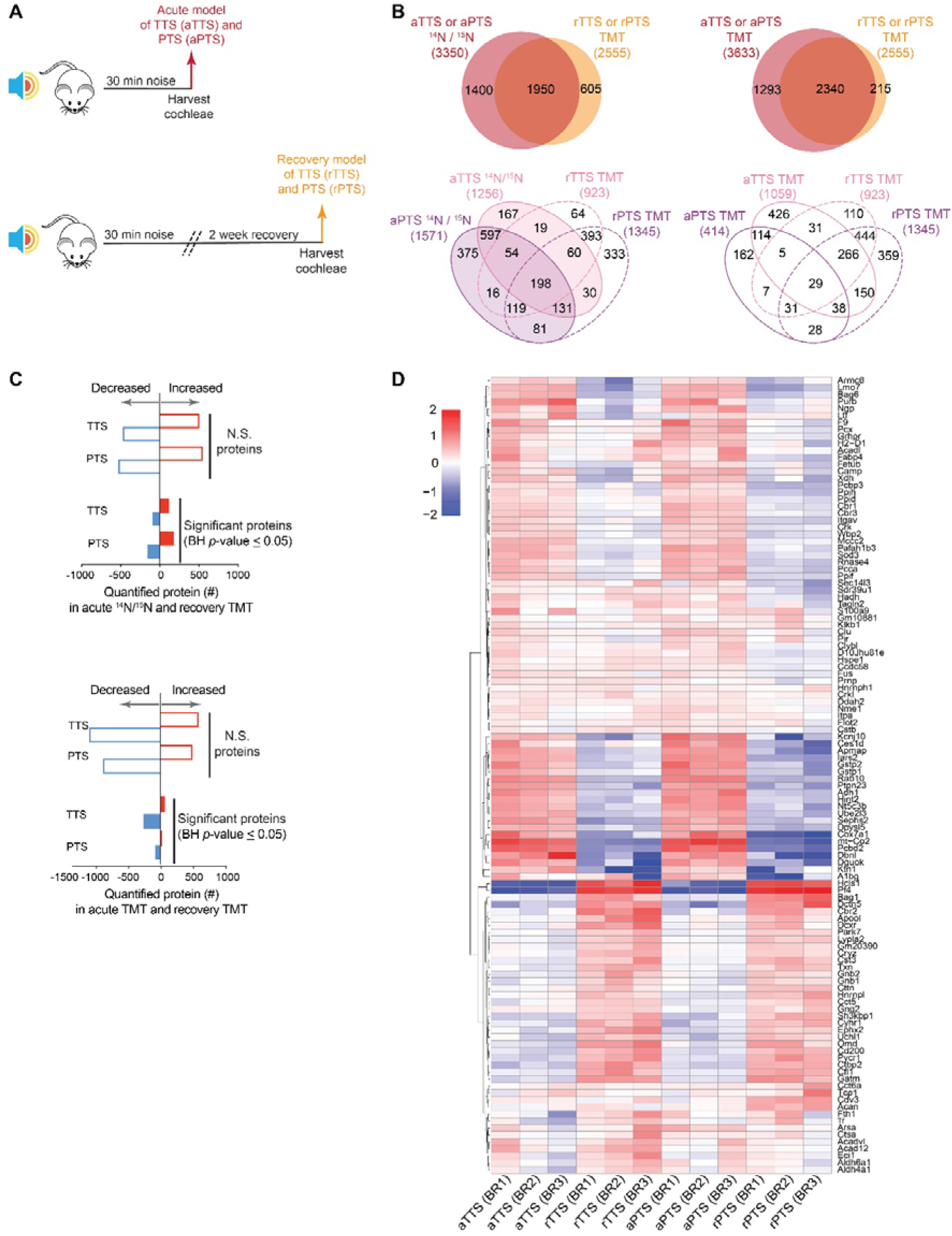
Quantitative analysis of the cochlear proteome after two weeks of recovery. (*A*) Experimental scheme for identifying and quantifying proteins in acute (aTTS, aPTS) or recovery (rTTS, rPTS) time points after 30 min of noise exposure. (*B*) Venn diagram showing common proteins measured in TTS and PTS conditions. (*C*) Number of quantified and significant protein verified according to their increase or decrease level of expression. (*D*) Heat map of significant protein across TTS and PTS conditions in acute and recovery time points.

Many proteins with increased levels after noise had reduced levels after recovery. Between the enriched Kyoto Encyclopedia of Genes and Genomes (KEGG) pathways and Gene Ontology (GO) molecular function associated with this dataset **(*SI Appendix*, Fig. S3)**, a majority of the acutely altered proteins were associated with metabolism (FDR = 1.16 × 10^-7^), catalytic activity (FDR = 2.2 × 10^-5^) and oxidoreductase activity (FDR = 3.02 × 10^-6^). More interestingly, the group of proteins with increased levels after the recovery period are involved with several biological process such as oxidation-reduction (FDR = 1.87 × 10^-4^), negative regulation of cell death (FDR = 2.69 × 10^-3^), and negative regulation of hydrogen peroxide-induced cell death (FDR = 2.84 × 10^-3^). Among the significantly altered protein in all four conditions at both time points were *Gstp1* and *Gstp 2* acutely and *Hcls 1*, *Park 7*, and *Gatm* after recovery. *Gstp 1* and *Gstp 2* are involved with detoxification via glutathione reduction (33). *Hcls 1* functions in the positive regulation on cell proliferation based on GO biological process (34). *Park 7* is a redox-sensitive chaperone, a sensor for oxidative stress, and protects neurons against oxidative stress (35). *Gatm* regulates cellular energy buffering and transport via creatine synthesis (36). Altogether, we identified a robust long-term cochlear proteostasis program in response to damaging levels of noise, which emphasizes protective cellular processes.

## Discussion

Nearly all previous attempts to determine the molecular mechanisms responsible for NIHL have been candidate-based approaches or have focused on changes at the mRNA level (37-39). These gene expression studies have provided important information regarding the underlying mechanisms of NIHL. We measured the level of more than 2,800 proteins in each noise exposure condition and revealed that the acute cochlear proteomic responses differ among three levels of noise intensity. Overall, the number of significant proteins identified increased with the intensity of noise. Although, our exposure levels were not expected to physically disrupt or damage individual polypeptides, we detected a large number of cytoskeletal proteins with consistently increased levels across higher noise intensities. Cytoskeletal proteins enriched in HCs have been widely studied in hearing research predominantly due to their important functions in HC stereocilia structure, cytoskeletal networks, and contractility of outer HCs (40-42). Increased levels of these proteins may suggest moderate to high levels of noise disrupt protein-protein interactions or drive a rapid reorganization of cytoskeletal protein complexes especially F-actin and myosin that were previously shown to have prompt responses within 10 min after calcium deprivation in cochlea (43) However, the degree of structural protein alterations may depend on multiple factors including the level and duration of exposure, for example, extended noise exposure induces actin depolymerization (44). Tubulin proteins are highly abundant in neurons and had increased levels after noise exposure. Presumably, due to ANFs swelling or other structural perturbations (45, 46). We also observed an increase in the levels of neurofilament proteins, which adds further support to our appreciation of ANFs as key noise substrates. Additional support for ANFs involvement comes from our GO analysis that identified several neuronal structures (synapse, neuron projection) especially at noise levels causing PTS. Interestingly, myelin associated proteins were enriched at all levels which supports previous reports that noise exposure may cause a loss of Schwann cells and contribute to permanent auditory deficits in NIHL (47). Septins, are actin and microtubule associated GTP binding proteins expressed by pillar and Deiter’s cells, and also efferent nerve terminals (48). Septins regulate in dendritic spine dynamics (49, 50), and collateral branching of axons (51). Sept 2,6,7,8,9 were significantly altered by all three levels of noise but Sept11 and Sept5, localize to cochlear efferent nerve synaptic vesicles were only significantly altered in TTS and PTS conditions (48). Septins have never been reported to be involved with NIHL. Increased levels of many cytoskeletal proteins immediately after noise exposure may reflect structural impairments or rapid reorganization and turnover.

Our observation that nearly the entire proteasome has increased levels after noise highlights the complex stress response triggered by noise exposure. We exposed mice for 15 or 30 minutes. These time frames are sufficient for rapid protein translation of existing mRNAs (52). Consistently, we identified many abundant proteins with elevated levels after noise. These proteins may be prominent in our datasets due to the fact that their mRNAs are highly abundant and selectively translated as part of the stress response (53). Gene transcription and posttranscriptional mechanisms could increase protein levels but it is unlikely since they likely require longer periods. Selective protein degradation by proteosomes, autophagosomes, and lysosomes are most likely to reduce the levels of distinct proteins. Extracellular proteins are likely to be degraded by additional proteases ***(SI Appendix)***.

STRING analysis revealed several important protein homeostasis regulatory networks that increased with higher levels of noise stimulation. For example, a large collection of proteasomal endopeptidases had increased levels after noise exposure intensities. As far as we are aware, there is no previous evidence that the proteasome is involved with NIHL. The proteasome is the major cellular degradation machine of the ubiquitin-proteasome system (54), is responsible for the bulk degradation of misfolded and damaged proteins (55), and is essential for cells to withstand and recover from various environmental stresses (56). Heat shock proteins (HSP) are also major noise sensitive substrates. Cells in the cochlea express HSPs after noise exposure and play protective roles (57, 58). Our results support and extend these findings, *Hspa 1a* (Hsp72), *Hspa 1b* (Hsp70) and the HSPs *Hsp 90aa 1* and *Hsp 90ab 1* had significantly increased levels both in TTS and PTS conditions. The third group of proteins with significantly increased according to noise stimulation are proteins that are involved in the mitochondrial electron transport chain (59). The complex I related protein, NADH: ubiquinone oxidoreductase or NADH dehydrogenase (*Nduf*) and complex III related protein, ubiquinol-cytochrome C reductase (*Uqcr*) had significantly increased levels in PTS and suggests it is a candidate for superoxide generation.

We used two different quantitative MS based methods to investigate changes in protein levels after TTS and PTS acoustic overexposure. Metabolic stable isotope labelling provides a very accurate and precise quantitative method both *in vitro* (60) and *in vivo* (61). Isobaric peptide labeling strategies are also powerful since up to twelve or more samples can be multiplexed and analyzed in the same MS analysis run (62). We used the TMT isobaric tags to confirm altered levels of hundreds of proteins from our metabolic labeling-based results. We did not reproduce the precise proteins and levels between the multiple datasets but the overall patterns and trends between the two strategies are in agreement. Similar to any other high throughput analysis method, MS has intrinsic technical challenges and biases. Confounding factors include biological sources such as animal to animal variation, experimental variations during sample processing, technical differences (63), and differences in the bioinformatic analysis. Altogether, these factors complicate our ability to directly compare global proteomic datasets.

We focused on noise exposures that induce “*auditory neuropathy*” which has been reported to deteriorate IHC synaptic compartments and produce functional decline based on ABR and DPOAE analyses (64). The criteria for synaptopathy-inducing TTS in a noise overstimulation model has been described in young adult mice (16 week, male CBA/CaJ) exposed to noise (8–16 kHz octave-band) at 100 dB SPL for 2 hours (10). However the pattern of hearing loss varies depending on differences in the age, sex, and strain of the mice (65, 66). While it is difficult to compare noise exposure between labs, due to variations according in the noise exposure chamber and loud speaker conditions, we show that neuropathy is also induced by shorter noise exposures and delineate conditions under which it occurs. More importantly, elucidation of the early protein biomarkers and substrates of different levels of noise exposure potentially provide new evidence regarding the molecular substrates of ANF afferent synapse damage.

We provide a pioneering proteomic description of both the acute response and recovery program after noise exposure causing TTS and PTS. More than 90% of the proteins quantified in the ^15^N datasets were also measured with TMT. The number of significantly altered proteins at the recovery time point was marginally reduced compared to the acute response. The majority of quantified significant proteins had increased levels in the acute response and decreased during the two week recovery period. Exposure to noise causing TTS causes a less dramatic proteome remodeling compared to PTS. Interestingly, there are a group of proteins with lowered levels immediately after noise but have increased levels during recovery. For example, Gstp1 and Gstp2, play an important role in detoxification by catalyzing the conjugation of hydrophobic and electrophilic biological molecules. An increase of glutathione related proteins levels correlates well with evidence in using glutathione to attenuate level of hearing deficit from noise exposure (67, 68). Analysis of the proteins involved in the recovery process after noise exposure highlighted several potential mitigators of noise-induced stress such as, Hcls 1, Park 7 and Gatm ***(SI Appendix)***. However, the vast majority of the proteins identified in the current investigation have never before been linked to acoustic injury. Therefore, future studies verifying their functional involvement in the regulation and prevention of the cochlear response to acoustic overstimulation are crucially important in providing new insights into the molecular basis of NIHL which will pave the path of therapeutic discovery in the near future.

## Acknowledgements

We thank Kwang Pak and Ann Hickox for their assistance. This work was supported by NIDCD/NIH (R00 DC-013805 to J.N.S.), NU Knowles Hearing Center, and VA (BLS Grant BX001295 to A.F.R.).

